# Single Image-based Vignetting Correction for Improving the Consistency of Neural Activity Analysis in 2-Photon Functional Microscopy

**DOI:** 10.1101/2021.03.01.433412

**Authors:** Dong Li, Guangyu Wang, René Werner, Hong Xie, Ji-Song Guan, Claus C. Hilgetag

**Affiliations:** Institute of Computational Neuroscience, University Medical Center Hamburg-Eppendorf, Hamburg, Germany; School of Life Science and Technology, ShanghaiTech University, Shanghai, China; Center for Biomedical Artificial Intelligence (bAIome), University Medical Center Hamburg-Eppendorf, Hamburg, Germany; Zhangjiang Laboratory, Shanghai Research Center for Brain Science and Brain-Inspired Intelligence, Instiute of Brain-Intelligence Technology, Shanghai, China; Institute of Psychology, Chinese Academy of Sciences, Beijing, China; Department of Health Sciences, Boston University, Boston, MA, United States

**Keywords:** vignetting correction, functional microscopic imaging, neural activity, image analysis, imaging artifacts

## Abstract

High-resolution functional 2-photon microscopy of neural activity is a cornerstone technique in current neuroscience, enabling, for instance, the image-based analysis of relations of the organization of local neuron populations and their temporal neural activity patterns. Interpreting local image intensity as a direct quantitative measure of neural activity presumes, however, a consistent within- and across-image relationship between the image intensity and neural activity, which may be subject to interference by illumination artifacts. In particular, the so-called vignetting artifact - the decrease of image intensity towards the edges of an image - is, at the moment, widely neglected in the context of functional microscopy analyses of neural activity, but potentially introduces a substantial center-periphery bias of derived functional measures. In the present report, we propose a straightforward protocol for single image-based vignetting correction. Using immediate-early-gene-based 2-photon microscopic neural image data of the mouse brain, we show the necessity of correcting both image brightness and contrast to improve within- and across-image intensity consistency and demonstrate the plausibility of the resulting functional data.

## 1 Introduction

Modern microscopic imaging techniques provide unique insights into the structure and functioning of complex biological neural systems. For many applications – ranging from whole-brain imaging^1^ to single-cell isolation^2^ – it has been shown that illumination correction, that is, the removal of uneven illumination of the scene and specimen, facilitates image interpretation, improves visibility particularly of fine structures and is a pivotal pre-processing step for subsequent image analysis^3^. Illumination correction is, therefore, often an essential component of microscopy imaging setups^4,5^ and image post-processing^6–8^.

A common and major illumination artifact in microscopic images is the *vignetting artifact*, which reflects a (usually radial) decrease of the image brightness, its contrast, or the saturation from the image center towards the periphery^9^. Correction approaches can be divided into prospective methods that exploit reference images^10^, retrospective multiple image-based correction^3,11^ and single image-based strategies. Reference and multiple image-based methods are considered most reliable^3^, but not always applicable due to application-specific constraints (e.g., data availability). At this point, single image correction strategies come into play^9,12^.

In the present report, we focus on high-resolution functional microscopic imaging and its application to studying the functioning of biological neural systems. Techniques such as calcium indicator-based^13–19^ or immediate early gene (IEG)-based imaging^20–24^ allow, in principle, an interpretation of the read-out of local image intensity as neural activity. While, intuitively, a quantitative comparison of neural activity of, e.g., local neuron populations is susceptible to illumination artifacts, illumination correction has, so far, been widely neglected or unreported in respective studies. A potential reason might be that established reference- and multiple image-based correction methods potentially affect the analysis of interrelations of neural activity patterns represented in different microscopic images, by inducing artificial correlations between the modified images. To overcome this issue,

- we develop and propose a straightforward single image-based vignetting correction protocol and
- demonstrate the value of the vignetting correction using single channel IEG microscopic images of the mouse brain.

Due to the application-inherent lack of ground truth data to evaluate the effect of the illumination correction, we focus on data plausibility: In the presence of a vignetting artifact, the functional data contain a significant center-periphery bias of neural activity measures that, for the performed experiments, cannot be accounted for by the underlying biology. Considering the reduction of the center-periphery bias in neural activity as well as consistency of absolute and relative activity changes over time as proxies of success, we demonstrate that the vignetting correction substantially improves data plausibility. Going beyond the standard approach of only taking into account the brightness of the image background^9^, we further show that the largest improvement in data plausibility is achieved by correction of both image brightness and contrast.

## 2 Materials and Methods

### 2.1 Imaging and Image Data

The present study is based on single channel 2-photon imaging mouse data as detailed in Xie et al.^23^ The mouse strain was BAC-EGR-1-EGFP (Tg(Egr1-EGFP)GO90Gsat/Mmucd from the Gensat project, distributed by Jackson Laboratories. Animal care was in accordance with the institutional guidelines of and the experimental protocol approved by Tsinghua University. To allow for *in vivo* imaging, a cranial window was implanted between the ears of 3-5 months old mice. Data recording started a month later. EGFP fluorescent intensity was imaged using an Olympus Fluoview 1200MPE with pre-chirp optics and a fast AOM mounted on an Olympus BX61WI upright microscope, coupled with a 2 mm working distance, 25x water immersion lens (numerical aperture 1.05). The animals were anesthetized 1 h after they explored a multisensory environment. Under the given experimental conditions, anesthetization is known to have very little effect on protein expression^25^, and protein expression to reflect neural activity.^23^ We randomly selected one image stack with measurements acquired at two days (interval in between: 5 days) to showcase our analysis. The stack belonged to the primary visual (VISp) area. Stack size was 512 × 512 pixels with a pixel edge length of 0.996 μm for the in-plane slices, containing 350 slices in z-direction with a spacing of 2 μm. Neuron segmentation was performed as described in Xie et al.^23^ The segmentation was performed in the uncorrected image data, to avoid neuron position mismatches between the unprocessed and the corrected image data. The neuron center positions were used to evaluate neural activity.

### 2.2 Illumination Correction Pipeline

Our proposed single image-based pipeline for vignetting correction in 2-photon functional microscopy data is outlined in **Figure**. Let a raw, i.e. measured and potentially post-processed single channel microscopic image be denoted by 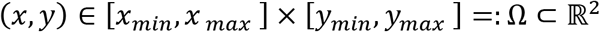. In the following, we approximate the relationship between *I*_0_ and the sought true image and neural activity *I*, respectively, by

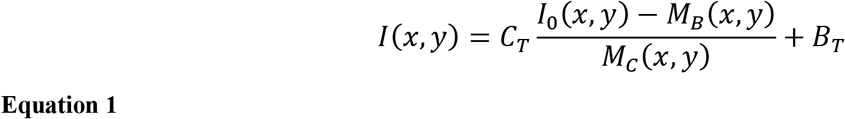

with *M_C_*(*x,y*) and *M_B_*(*x,y*) as a spatially varying gain or contrast distribution and a spatially varying background brightness, respectively, that are to be estimated and compensated during the illumination correction. *B_T_* and *C_T_* are pre-defined additive and multiplicative constants to bring the corrected image into a desired value range^26^, and are not part of the correction process. With *C_T_* = 1 and *B_T_* = 0, equation 1 corresponds to the standard notation as, for instance, used by Smith et al.^3^

The proposed correction approach follows the common assumption that vignetting effects can be approximated by Gaussian functions. Leong et al.^9^, for instance, described the uneven illumination as an additive low frequency signal and approximated it by an isotropic Gaussian distribution with large standard deviation. The distribution parameters were, however, chosen ad-hoc and image-specific.^9^ Here, we also model *M_B_*(*x,y*) and *M_C_*(*x,y*) by 2-dimensional (2D) Gaussian functions, but estimate the distribution parameters by analysis of *I*_0_(*x, y*).

To be able to efficiently cope with potentially large image data sets, we employ a patched-based approach, consisting of two main steps:

[STEP 1] patch-based robust estimation of local background brightness and contrast for a sufficient amount of supporting points distributed across *I*_0_(*x,y*), and
[STEP 2] estimation of *M_B_*(*x,y*) and *M_C_*(*x,y*) based on the supporting point brightness and contrast values.

#### 2.2.1 STEP 1: Patch-based Estimation of Local Brightness and Contrast

In line with most existing methods in the given context, we focus on the image background to estimate the vignetting functions.^9,27,28^ Let 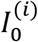 denote a patch of *I*_0_ with a patch center (*x*^(*i*)^, *y*^(*i*)^) ∈ Ω and a domain [*x_i_* – *λ_x_*/2, *x_i_* + *λ_x_*/2] × [*y_i_* – *λ_y_*/2, + *λ_y_*/2] =: Ω_i_ ⊂ Ω, with *λ_x_* and *λ_y_* as the side lengths of the patches. Subsequently, we assume an ordered sequence of non-overlapping patches 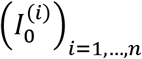 to be given that covers *I*_0_. However, the proposed approach is applicable to any patch sampling strategy. The patch centers are considered the supporting points for [STEP 2]; the associated brightness and contrast values were computed based on a histogram analysis of the intensity values of the patch pixels. Starting with the original histogram for a patch 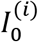, contributions by high intensity, that is, foreground objects such as highly active neurons that lead to a long-tailed intensity distribution, were removed following the approach of Alstott et al.^29^and Clauset et al.^30^ that seeks to find the minimum histogram data point to optimally fit a power law to the right tail of pixel intensity distribution. The histogram data points above this intensity value were discarded from further analysis. Further remaining high and low intensity values were removed by focusing on the central parts of the histogram and the intensity distribution, respectively. The intensity distribution of the remaining data points was tested for normality using the Shapiro-Wilk test^31^; only patches with a resulting value above a predefined threshold of the test statistic were considered valid supporting points for [STEP 2]. The corresponding local brightness and contrast values *B*(*x*^(*i*)^, *y*^(*i*)^) and *C*(*x*^(*i*)^, *y*^(*i*)^) were approximated by the expectation and the standard deviation of the fitted normal distribution. For further details and parameter values used in our study, we refer to the source code (see Data Availability).

#### 2.2.2 STEP 2: Approximation of *M_B_*(*x,y*) and *M_C_*(*x,y*)

Based on the patches 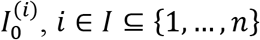, for which the intensity distributions passed the Shapiro-Wilk test, the corresponding set of patch estimates *B*(*x*^(*i*)^, *y*^(*i*)^) and *C*(*x*^(*i*)^, *y*^(*i*)^) were used to fit 2D-Gaussian distributions to estimate the sought distributions *M_B_*(*x,y*) and *M_C_*(*x,y*).

### 2.3 Experiments and Evaluation

To evaluate plausibility of the image correction, the slices of the image stack (see Sec. 2.1) were evenly divided into 4 × 4 = 16 regions (see Fig. 2, panels A-D): four center regions (C1 to C4), eight edge regions (E1 to E8), and four angle regions (A1 to A4). If a vignetting effect exists, the comparison of neuronal activity derived for the different regions shows a significant bias, by which the center regions have high activity, which decreases radially (i.e., highest activity differences between the C and the A regions). Correspondingly, after successful correction of the vignetting effect, the bias should be significantly reduced. The slices were further grouped into four laminar compartments: layer II/III slices (95 slices in our selected stack), layer IV slices (41 slices), layer V slices (94 slices), and layer VI slices (97 slices). The layers were assigned manually.

Motivated by functional neuroimaging studies^13–25,32^, we considered the following measures to analyze the impact of the vignetting effect and the reduction thereof:

- **Neural activity:** Based on the segmentation of the neurons (see Sec. 2.1), the activity *X_l_* of the *l*-th detected neuron (*l* = 1,…, *n_l_*) is computed as the intensity of the center pixel of the segmentation mask.
- **Absolute changes of neural activity over time:** Neural activity measurements were available for two different days, denoted as *day1* and *day0*, with an intervening interval of five days. In addition to the neural activity, we also evaluated the absolute activity changes Δ*X_l_* = *X_l_*(*day1*) – *X_l_*(*day0*).
- **Relative changes of neural activity:** In addition, we evaluated the relative neural activity changes *δX_l_* = [*X_l_*(*day1*) – *X_l_*(*day0*)]/[*X_l_*(*day1*) + *X_l_*(*day0*)].

The three measures *X_l_*, Δ*X_l_* and *δX_l_* were separately evaluated for the 16 regions (C1-C4, E1-E8, A1-A4) and the four laminar compartments (II/III, IV, V, VI) before and after vignetting correction. Results are given as mean and standard deviation (STD) of the measures for the individual regions and laminar compartments. Significant differences between the measures of two different regions were evaluated separately for the different laminar compartments, applying t-tests with Bonferroni correction of the *p*-values. Moreover, for each pair of regions, the relative difference *δ_STD_* of the standard deviations of the considered measure values within the individual regions was computed for the different laminar compartments. Thus, for each laminar compartment, in total *n_c_* = 6 centre-centre (C-C) comparisons, *n_c_* = 28 edge-edge (E-E) comparisons, *n_c_* = 6 angle-angle (A-A) comparisons, *n_c_* = 32 centre-edge (C-E) comparisons, *n_c_* = 32 edge-angle (E-A) comparisons, and *n_c_* = 16 center-angle (C-A) comparisons were performed.

To obtain further insights into a potential vignetting-associated center-to-periphery bias before image correction, we also evaluated the fraction of region pairs with non-significant differences (*p* ≥ 0.01 after Bonferroni correction) for the different measures on a region-type level (i.e., focusing on all CC, E-E, A-A, C-A, C-E, etc. comparisons). Let *c_p_* = *n*_*c*(*p*≥0.01)_/*n_c_* denote the fraction of the number specific region type-specific comparisons (e.g. *n_c_* = 16 for C-A comparisons) and the number *n*_*c*(*p*≥0.01)_ of corresponding non-significant comparisons. Then, the ratio Δ*C* = *c_p_*/〈*c_p_*〉_*same*_ was computed, with 〈*c_p_*〉_*same*_ as the average value of *c_p_* values of the C-C, E-E, and A-A comparisons. The hypothesis was that, in the presence of a vignetting effect, Δ*C* values for, e.g., the C-E and C-A comparisons are considerably smaller than one; after successful correction, the values should become closer to one. In particular, we assumed the comparison of Δ*C* values for C-A and E-E comparisons before and after correction to indicate presence and successful correction of the vignetting effect.

Similar to Δ*C*, we also evaluated differences of the relative standard deviations *δ_STD_* on a region-type level. With 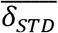 as average *δ_STD_* value for all pairs of two of the region types C, E or A for a specific layer compartment, we computed 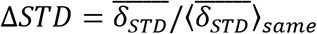, with 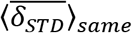 denoting the average value of 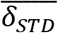 for C-C, E-E, and A-A comparisons. A correction of a vignetting effect would, for instance, lead to significant reduction of Δ*STD* for C-A comparisons, especially when compared to Δ*STD* for E-E comparisons.

All measures were evaluated for the original image data as well as after vignetting correction according to equation 1. In addition, we also applied a brightness-only correction, using

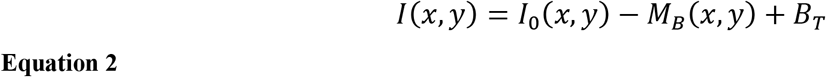

as well as a contrast-only correction,

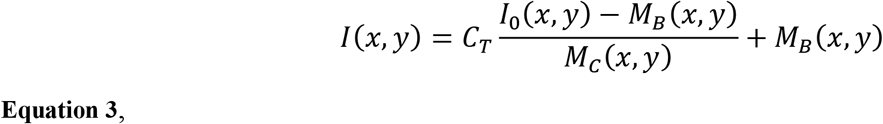

to illustrate the respective effects.

## 3 Results

Subsequent results are based on automatically segmented 14.662 neurons. Within each region of the laminar compartments, on average, 229 (STD: 78; range: 50 – 376) neurons were segmented. An example of an image slice before and after vignetting correction is shown in Figure 1, Panels B and H, respectively. In the original data, a strong image intensity decrease from the center toward the periphery can be seen. This gradient, in turn, leads to a larger number of significant differences of the measures derived for the different regions and region-types, indicated by blue and green fields in the lower triangles of Panel E of Figure 2 (focusing on neural activity *X_l_* itself), Figure S1 (absolute changes Δ*X_l_* of the neural activity), and Figure S2 (relative changes *δX_l_* of neural activity), respectively, for the layers II/III of the considered VISp area. Corresponding representations for layers IV, V and VI are shown in Figs. S3-S17 of the supplementary data. Analogously, large *δ_STD_* values are present for the original data (see lower triangle of Panel F of Fig. 1 and Figs. S1-S17). In particular, the C-A comparisons exhibit a larger number of significant differences and larger *δ_STD_* values than the “within-type” comparisons, that is, the C-C, E-E and A-A comparisons. Corresponding C-E and E-A values are somewhere in between. This finding clearly indicates the existence of vignetting effects in the original data.

**Figure 1.**
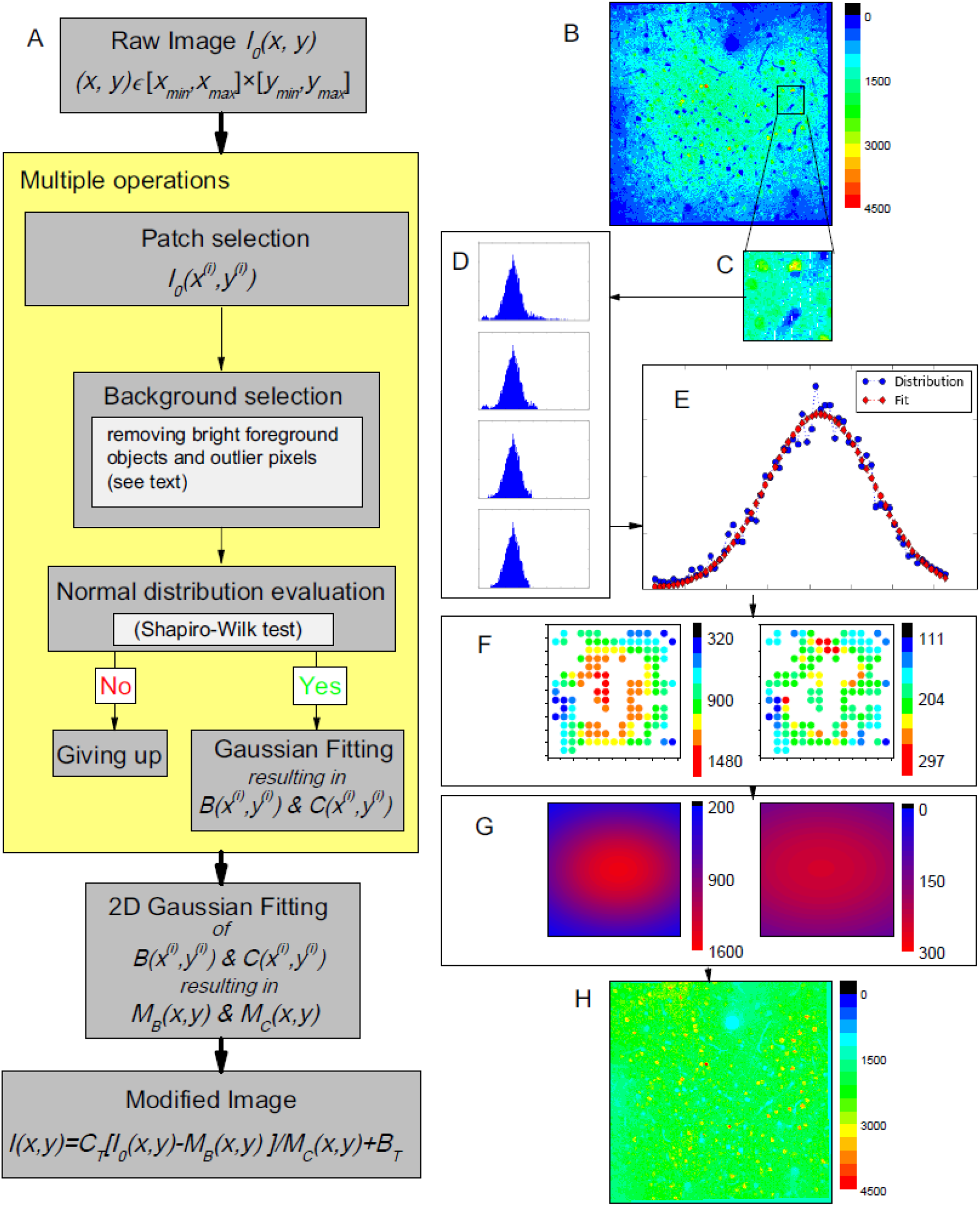
(A) Working pipeline. (B-H) One example of the proposed vignetting correction of a 2-photon microscopy image of mouse VISp area at layers II/III depth (200 μm depth). (B) The original image, i.e. before correction. (C) Exemplary patch selection. (D) Histograms of image pixel intensities, illustrating the effect of the performed outlier reduction. In top-down direction: first, histograms of all pixels in the patch; second, after dropping off the long tail; third, after dropping off some data of certain values at two ends; last, after dropping off additional data of certain amounts at two ends (details given in the source code). At the end, the similarity to a normal distribution estimated by a Shapiro-Wilk test is 0.996, larger than the threshold we used (0.98). (E) Corresponding Gaussian fitting of the background intensity for the patch pixels. (F) Supporting point (i.e. the patch centers) brightness *B*(*x*^(*i*)^, *y*^(*i*)^) (left panel) and contrast *C*(*x*^(*i*)^, *y*^(*i*)^) (right panel). (G) The estimated global brightness distributions *M_B_*(*x,y*) and *M_C_*(*x,y*) (left and right panel). (H) The final modified image.

**Figure 2.**
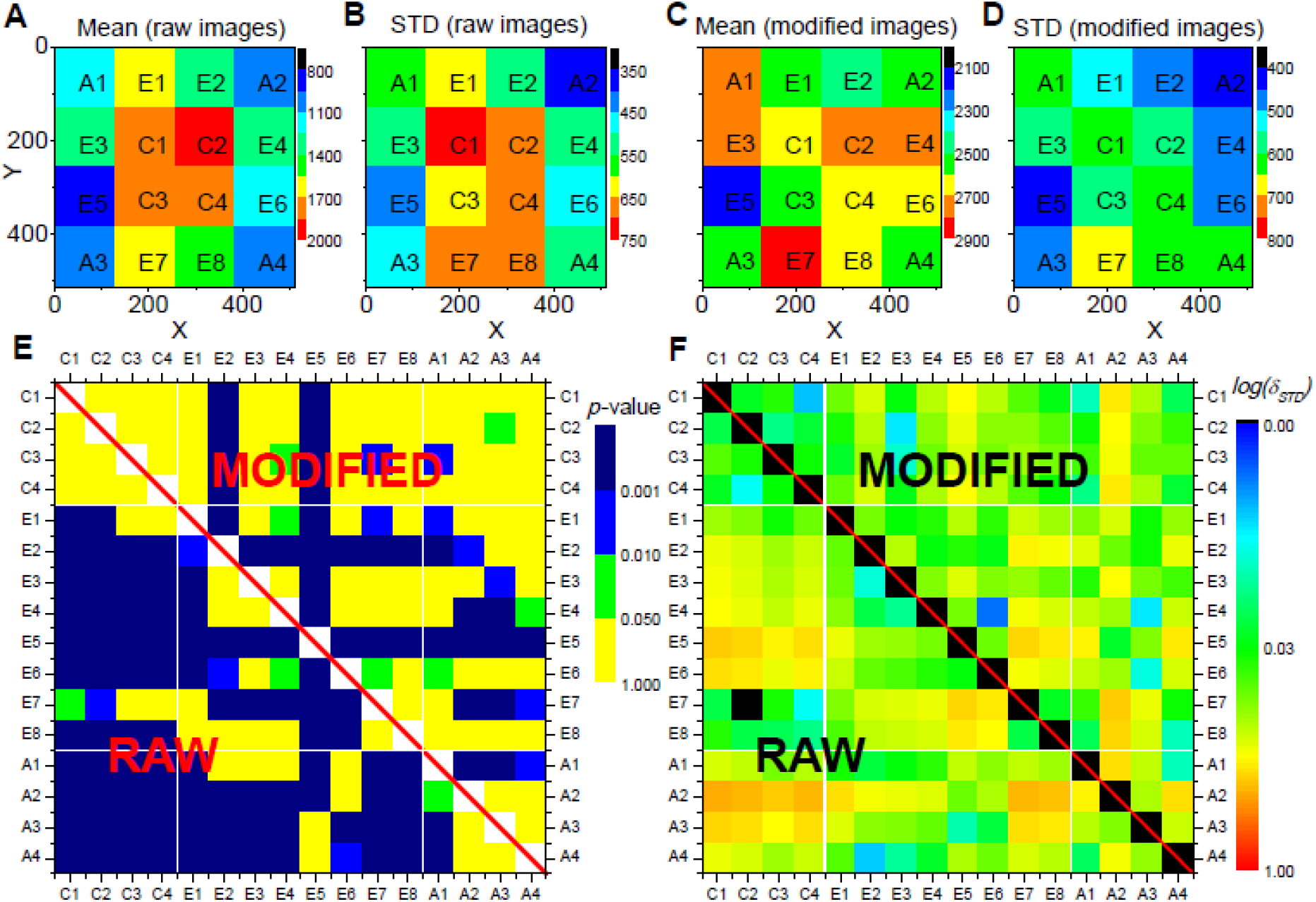
Evaluation results for the proposed vignetting correction for the neural activities *X_l_* in layers II/III of a mouse VISp area. The mouse was located in homecage. The images stack was evenly divided into 16 regions on x-y plane, as shown in panels (A-D). Panel (A) shows the mean activities of all neurons within each region sampled from the raw images, and panel (B) shows their standard deviations. Panels (C) and (D) show the mean values and the standard deviations sampled from the modified images. Panel (E) shows the comparison of the neural activities between each pair of regions, where colors stand for the p-values of a respective t-test with Bonferroni correction. Panel (F) shows the comparison of the distributions of the neural activities between each pair of regions, where colors indicate δ_STD_ with logarithm scale. In Panels (E) and (F), the lower rectangles, i.e. the values below the diagonal, shows the comparison of the data from the original images. The upper triangles represent corresponding data from the modified images. All parameters for the vignetting correction are the same as applied in Fig. 1.

After applying the proposed image correction procedure, the abovementioned effects are substantially diminished: The number of significant region-to-region differences of the evaluated measures is considerably reduced; similarly, the *δ_STD_* values are more consistent between the regions of different type (e.g., C-A and E-A; see upper triangle in Panels E and F in Fig. 1 and Figs. S1-S11). This effect, however, is much less pronounced if only an intensity or only a contrast correction is applied (Figs. S12-S17).

In line with these observations and the hypothesis described in Sec. 2.3, before the vignetting effect correction, the Δ*C* values for the C-A comparisons are significantly lower (*p* < 0.01) and the Δ*STD* values significantly higher than corresponding values for E-E comparisons (*p* < 0.001). After image correction, the differences were no longer significant, associated with a significant increase of Δ*C* and decrease of Δ*STD* (both *p* < 0.001), due to the C-A values after image modification (Fig. 3 A and B).

**Figure 3.**
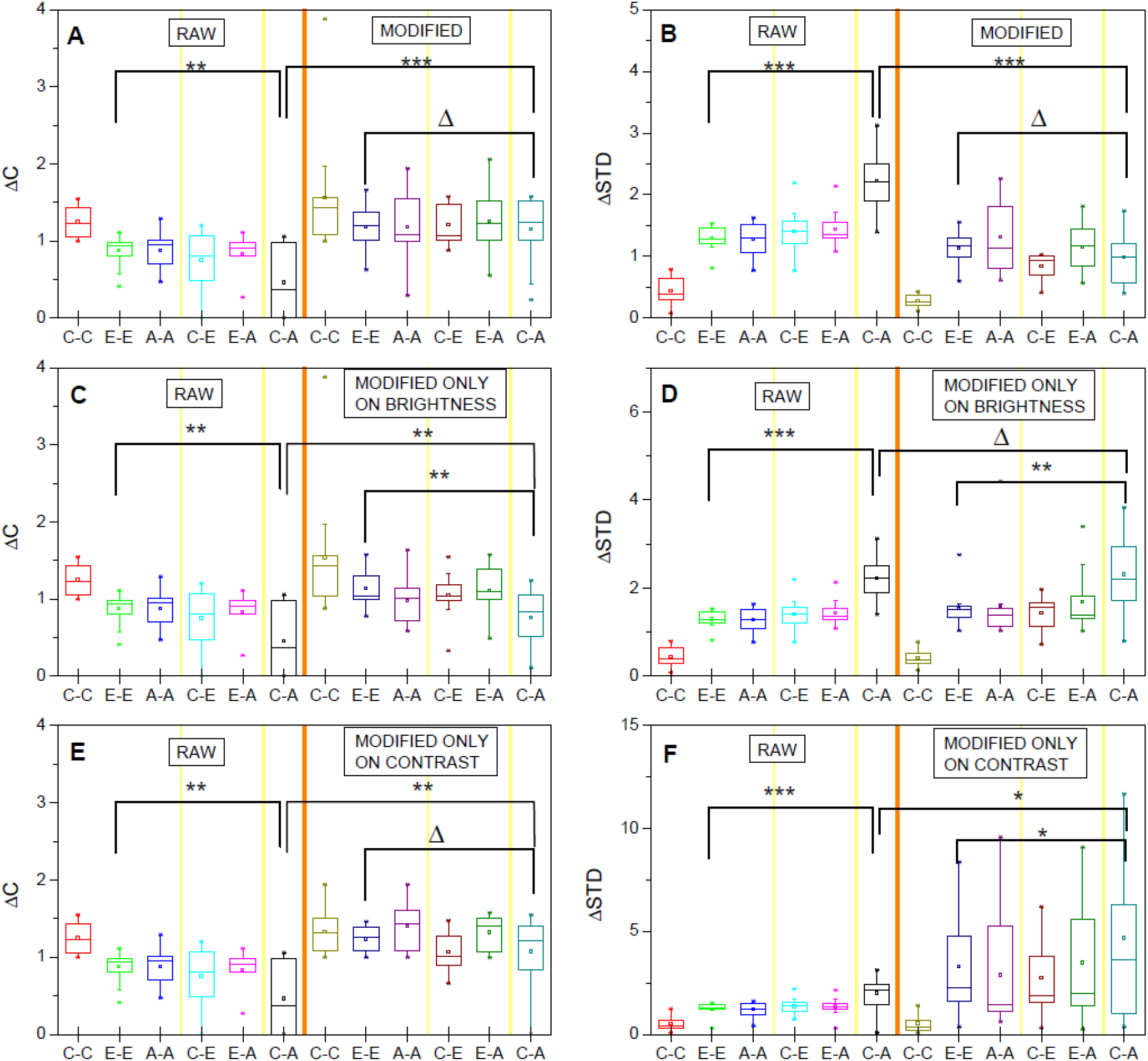
ΔC and ΔSTD values for the original (RAW) and corresponding values after vignetting correction. Panels A and B represent the effect if performing brightness and contrast correction. In panels C and D (right sides), only brightness was corrected. Similarly, panels E and F show the data for contrast-only correction. The shown data are averaged over the obtained normalized data for the three studied measures (neural activity, absolute differences of neural activity, relative differences) and the different layer compartments. Δ, p>0.05, *, p<0.05; **, p<0.01, ***, p<0.001, t-test.

Fig. 3 C-F further reveal (similar to Figs. S12-S17) that brightness-only or contrast-only correction does not sufficiently correct for the vignetting effect. If only a brightness correction is applied, Δ*C* values are, for instance, still significantly lower for C-A comparisons than for E-E comparisons. If only a contrast correction is applied, the C-A Δ*STD* values are still significantly higher compared to the EE values (*p* < 0.05).

## 4 Discussion

### 4.1 Center-periphery bias and data rationality

Correction of uneven illumination, and in particular a correction of the vignetting effect, has become a standard procedure in *structural* neural microscopy image processing^1,2^. In contrast, the application of such corrections to corresponding *functional* neural microscopy images is rarely reported, and the influence and effects of respective procedures remain unclear so far. This lack of studies may, in part, be due to a lack of ground truth data that could be used for evaluation purposes. However, we hypothesize that the intrinsic consistency and plausibility of the data is an appropriate indicator for the existence and the success of a correction of vignetting effects. While neural microscopy data and commonly derived measures such as neural activity and absolute or relative activity changes over time certainly exhibit spatially varying distributions, there is no convincing physiological explanation for the existence of a distribution of the measures with a center-periphery bias–that is, a vignetting effect. A reduction, and ideally the elimination, of a center-periphery bias can, therefore, be studied to assess the success of a correction in functional microscopic neural images.

In this work, we implemented a straightforward single image-based correction method, and used IEG 2-photon microscopic neural image data of the mouse brain to demonstrate that such data are affected by vignetting effects (see Figure 1 B for the original image data and Panels A and of **Figure** and the Supplementary Figs. S1-S17 for derived measures), and that the proposed method corrects for the resulting center-periphery bias of the image intensity and derived measures (see, e.g., Panels C and D of **Figure**). We further illustrated that the most effective correction of the vignetting effect requires correction of both the brightness and the contrast of the image background. The corresponding code is publicly available to allow re-use of the proposed solutions (see Data Availability statement).

### 4.2 Multiple image-vs single image-based correction

As described in the introduction, vignetting correction approaches can be divided into two main groups: multiple image-based and single image-based correction. A prerequisite of multiple image-based vignetting correction is, obviously, the availability of multiple images. Multiple image correction is often considered more reliable than single image correction, but also comes with some potential disadvantages. One aspect is that multiple image-based correction typically exploits the intensity distribution of pixels at the same spatial position but in different images. This relationship, in turn, implicitly assumes and induces a (potentially artificial) correlation between the images that are processed. The artificial correlation may not be of utmost relevance when processing structural microscopy data. In functional microscopy neural data analysis, however, existence or absence of a correlation of signals from different images is often at the core of the studied research question^23,24^. Thus, it has to be ensured that the observed correlations are of a biological nature, rather than artificial.

This requirement, in turn, renders single image-based image correction particularly useful for functional microscopic neural data analysis; it is applicable even if the exact form of the vignetting effects varies day-by-day. For functional neural microscopy imaging, such day-by-day variations exist due to several reasons. For instance, one such reason is that blood vessel volume and throughput can vary, with a substantial impact on light intensity^33^. It is, in principle, technically possible to modify the craniofacial light intensities to a certain degree by adjusting the laser parameters, but this adjustment only eliminates the effects caused by vessels over the brain surface. Since there are far more unpredictable components influencing the light intensities inside the brain, e.g., the irregular distribution of blood vessels and dynamic locations of glia or immune cells, it is hardly possible to preset proper parameters at different depths to unify the intensities across days for both the superficial and deeper slices at the same time. In addition, daily shifts of the position of the mouse brain in the microscope induce additional sources of error. The errors are corrected by slightly rotating the images after scanning, but this inevitably also changes the appearance of the vignetting effects. Moreover, the optical pathways of the laser cannot be guaranteed to be sufficiently stable on long time scales. If the experiments last too long, this may also be a reason for daily variations of observed vignetting effects.

Furthermore, from a methodical perspective, underlying mechanisms of multiple and single image-based correction approaches are often, to a certain degree, similar. For example, Smith et al.^3^ simultaneously modified image brightness and contrast, similar to our proposed approach. For both approaches, the image modification is based on the analysis and transformation of a pixel intensity distribution – with the difference that Smith et al.^3^ determined the distribution of pixels from different images, but at the same spatial position. In contrast, we focus on patch pixels of a single image; in addition, we exclude bright objects (here: mainly pixels of high activity) to ensure an unbiased estimation of the background intensity distributions.

Thus, for single image-based vignetting correction, the multiple image-based prior information is essentially replaced by the hypothesis that the image background intensity and the image contrast distribution can be approximated by two-dimensional Gaussian distributions. The two-dimensional Gaussian fitting to the image background for single image-based vignetting correction was already proposed by Leong et al.^9^. In the present work, their idea was extended by additional consideration of the image background contrast. The described results indicated that both brightness and contrast correction appear essential. In addition, high R^2^ values for the fitting reveal appropriateness of the underlying assumption (R^2^ values of 0.9 and 0.75 for brightness and contrast fitting, respectively, for more than approximately 170 valid patches; see Supplementary Fig. S18).

### 4.3 Unifying the brightness and contrast in superior-inferior direction

The constants *B_T_* and *C_T_* can, in principle, be chosen arbitrarily and can be used to unify the mean background intensity and contrast across slices of a three-dimensional image stack, i.e., along the superior-inferior direction, which is also considered important for analysis of respective data^34,35^. In our experiments, we indeed did so and set the constants to identical values for all slices. It should, however, be noted that one should be careful not to over-interpret the actual intensity levels in terms of neuroscientific conclusions, e.g., by comparison of absolute intensity values across cortical layers. Similar to the explanation of the existence of potential daily variations of the vignetting effect, the amount of synapses, glia cells, vessels, etc. and, thus, the light intensity for the different slices will also vary. In turn, due the similar cellular composition for slices within a specific layer compartment, it appears reasonable to compare the intensity values within a laminar compartment. For a reliable comparison across laminar compartment borders, there is, however, a lack of supporting evidence.

## 7 Author Contributions

J-SG and CCH designed the research; DL and GW initialized the illumination correction idea; DL and RW developed the illumination correction and evaluation methodology; GW and HX worked on the animal experiments and data preparation; DL, RW, J-SG and CCH wrote the manuscript; and all the authors approved manuscript.

## 9 Data availability

The source code and datasets for this study will be made publicly available on github after acceptance for publication.

